# Vps4 substrate binding and coupled mechanisms of Vps4p substrate recruitment and release from autoinhibition

**DOI:** 10.1101/2024.09.07.611824

**Authors:** HJ Wienkers, H Han, FG Whitby, CP Hill

## Abstract

The ESCRT pathway’s AAA+ ATPase, Vps4p, remodels ESCRT-III complexes to drive membrane fission. Here, we use peptide binding assays to further the understanding of substrate specificity and the mechanism of autoinhibition. Our results reveal unexpected sequence preference to the substrate binding groove and an elegant mechanism of regulation that couples localization to substrate with release from autoinhibition.

## Main Text

The Vacuolar Protein Sorting 4 protein (Vps4p in *S. cerevisiae*) is an essential component of the Endosomal Sorting Complex Required for Transport (ESCRT) pathways that drive membrane fission in many biological processes, including cytokinetic abscission^1^, multivesicular body formation^2^, exosome release^3^, and virus budding^4,5^. The ESCRT pathway and Vps4p are conserved throughout eukaryotes, with Vps4 sharing ∼60% sequence identity with its two human homologs, VPS4A and VPS4B. Vps4p is thought to trigger membrane fission by unfolding ESCRT-III subunits, thereby remodeling ESCRT-III filaments that bind and stabilize highly distorted configurations of the lipid bilayer^6^. Because Vps4p appears to have potential to unfold a wide variety of proteins, it is critically important that its access to authentic substrates is tightly controlled.

Vps4p is a member of the meiotic clade of AAA+ ATPases^7^. Its structure comprises an N-terminal Microtubule-Interacting and Trafficking (MIT) domain followed by a flexible 29-residue linker and an ATPase cassette. The MIT domain is a three-helix bundle that mediates localization to substrate by binding MIT Interacting Motifs (MIMs) located near the C-termini of ESCRT-III subunits^8–10^. Multiple cryo-EM structures^11–15^, including complexes with an 8-residue peptide (peptide-8) derived from the ESCRT-III family member Vps2p^11,14,15^, show five of the hexameric Vps4p subunits forming a helix/spiral while the sixth subunit is found in transition between ends of the spiral. Within the spiral there are four subunit interfaces that each create Class 1 and Class 2 peptide binding pockets that are largely formed by the pore loop 1 residues W206 and M207, respectively. The bound peptide-8 adopts a β-strand conformation, with the side chains of the eight residues occupying the four Class 1 and four Class 2 peptide binding pockets (Figs. 1ab).

**Fig. 1.**
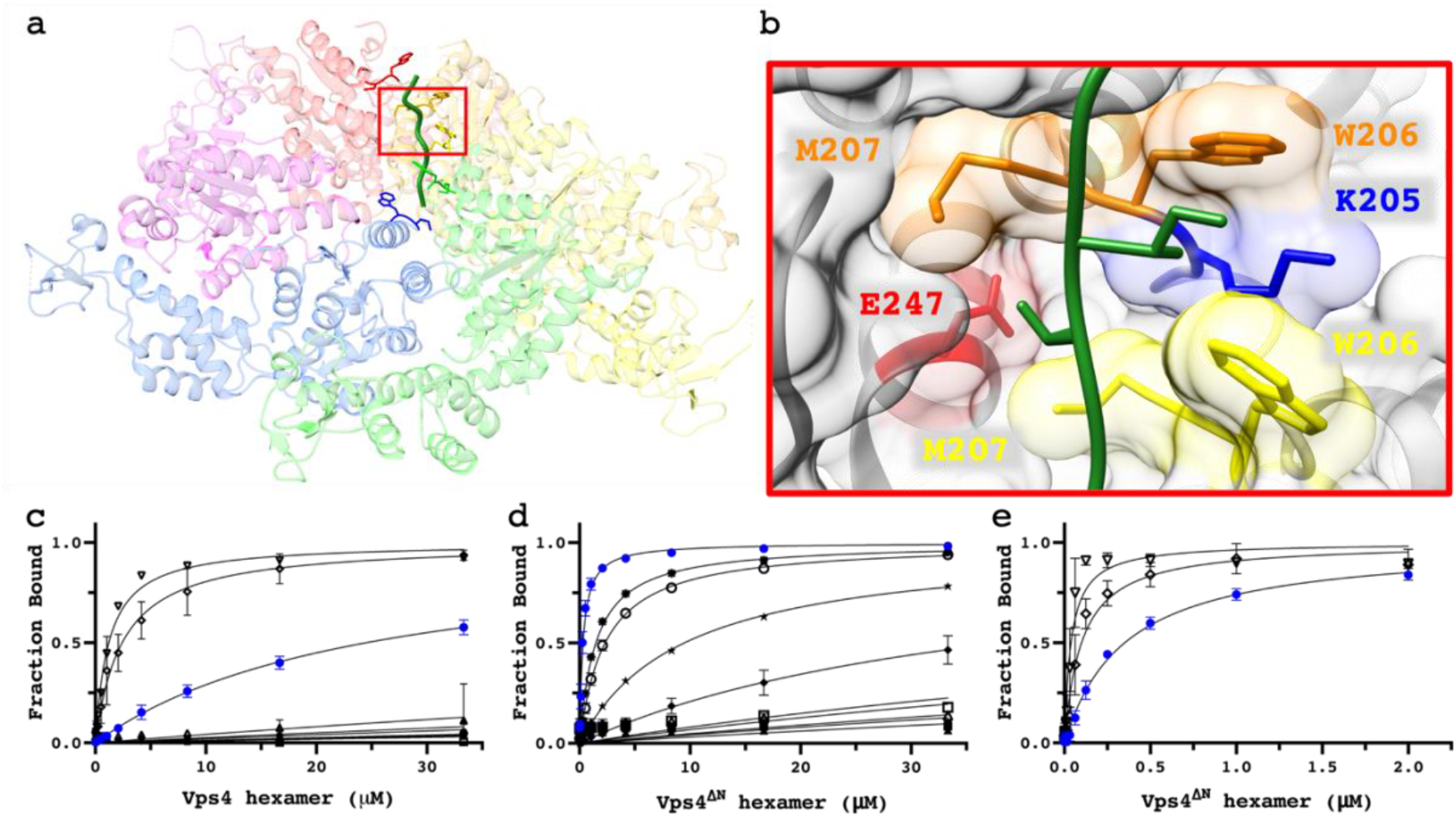
Binding to the translocation groove. **a**, Peptide-8 bound to the Vps4p translocation groove (PDB: 6AP1)^15^. **b**, Close-up of class 1 and 2 pockets created between W206 (orange and yellow) M207 (orange and yellow) respectively with charged contributions from K205 (blue) and E247 (red) respectively. **cde**, Fluorescence polarization peptide binding assays to Vps4p and Vps4p^ΔN^, see Table 1 for peptide associated symbols.

The structures have prompted a mechanistic model (Fig. 1a) in which hydrolysis occurs at the lowest subunit interface in the spiral to trigger disengagement of the lowest subunit from its neighboring subunit and the substrate^11^. Upon disengagement, the nucleotide-binding site is opened and can exchange ADP for ATP, which promotes reengagement at the top of the spiral. Repeated cycles will result in Vps4p “walking” along ESCRT-III, thereby forcing the substrate into an extended/unfolded conformation. Structural similarity to multiple other AAA+ unfoldases suggests that this mechanism is widely conserved^16,17^.

**Table 1.**
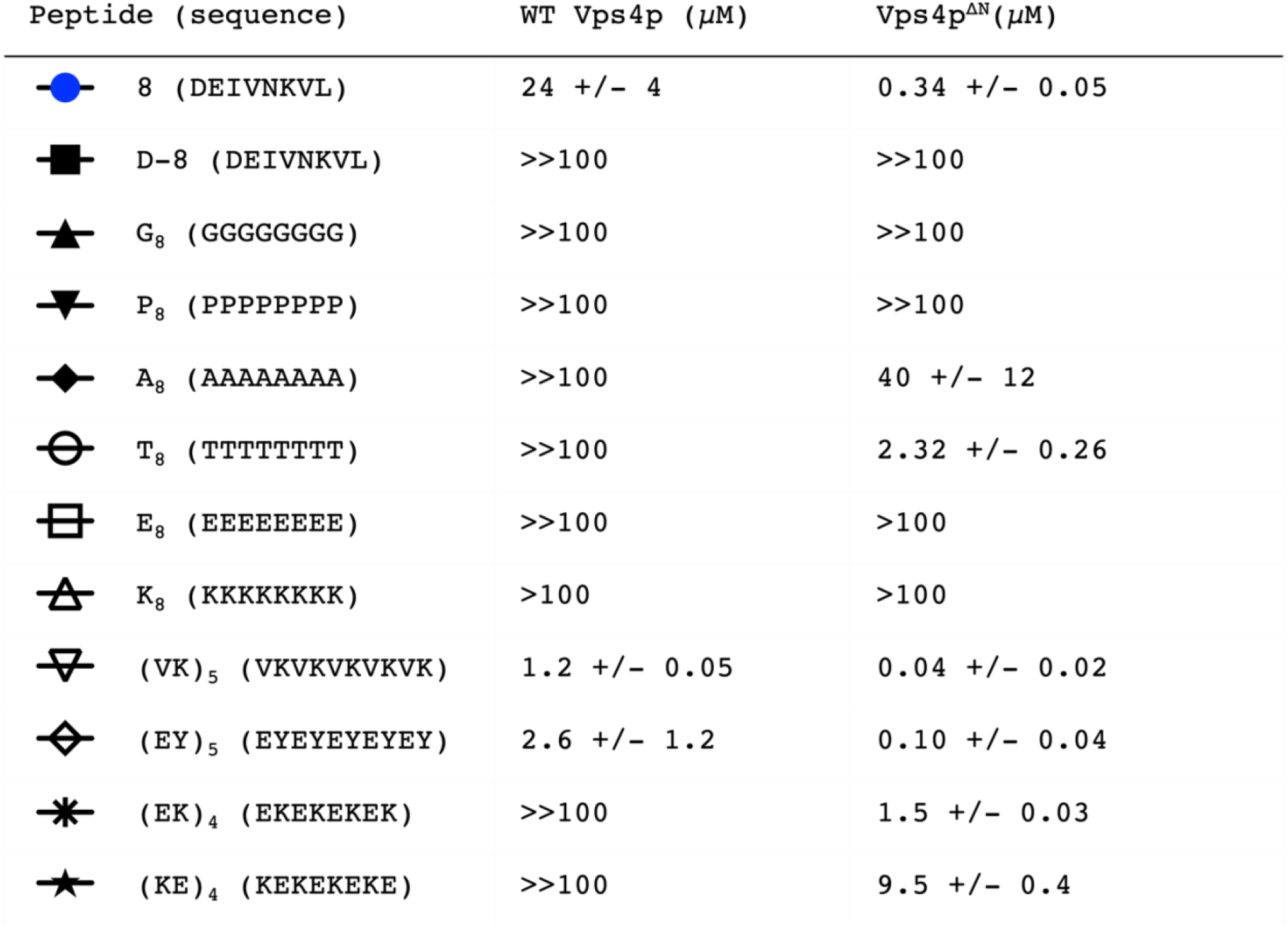
Peptide binding affinities of Vps4p and Vps4p^ΔN^. Data are shown in Figs 1d-f. Affinity is denoted >>100 μM when the fraction bound is below 10% at 33 μM Vps4p-hexamer, and >100 μM when the fraction bound is at or above 10% at 33 μM Vps4p hexamer but cannot be fit with confidence.

| Peptide (sequence) | WT Vps4p ($\mu\text{M}$) | Vps4p<sup>ΔN</sup>($\mu\text{M}$) |
| --- | --- | --- |
| 8 (DEIVNKVL) | 24 +/- 4 | 0.34 +/- 0.05 |
| D-8 (DEIVNKVL) | >>100 | >>100 |
| G<sub>8</sub> (GGGGGGGG) | >>100 | >>100 |
| P<sub>8</sub> (PPPPPPPP) | >>100 | >>100 |
| A<sub>8</sub> (AAAAAAAA) | >>100 | 40 +/- 12 |
| T<sub>8</sub> (TTTTTTTT) | >>100 | 2.32 +/- 0.26 |
| E<sub>8</sub> (EEEEEEEE) | >>100 | >100 |
| K<sub>8</sub> (KKKKKKKK) | >100 | >100 |
| (VK)<sub>5</sub> (VKVKVKVKVK) | 1.2 +/- 0.05 | 0.04 +/- 0.02 |
| (EY)<sub>5</sub> (EY EY EY EY EY) | 2.6 +/- 1.2 | 0.10 +/- 0.04 |
| (EK)<sub>4</sub> (EKEKEKEK) | >>100 | 1.5 +/- 0.03 |
| (KE)<sub>4</sub> (KEKEKEKE) | >>100 | 9.5 +/- 0.4 |

Consistent with the observation that peptide binds the Vps4p translocation groove in an extended beta conformation, peptide P_8_ (unable to form a beta strand), peptide G_8_ (lacking side chains), and the enantiomeric D-peptide-8 (mirror image rather than standard L-configuration) all showed weak binding against full length Vps4p (Table 1, Figs 1cde). They also showed weak binding affinity against the Vps4p^ΔN^ construct, which lacks the MIT domain and linker (residues 101-437) and, we found, displays higher peptide binding affinity than the full-length protein in an earlier study^18^. The conclusion that Vps4p binds peptides in a beta conformation leaves considerable potential to bind and translocate a wide variety of diverse sequences.

To further explore substrate binding, we determined the binding affinities of peptides: A_8_, T_8_, E_8_, K_8_, (VK)_5_, and (EY)_5_. While A_8_, T_8_, (VK)_5_, and (EY)_5_ showed appreciable binding affinity to Vps4p^ΔN^, Vps4p displays a preference for hydrophobic side chains (Table 1, Figs 1cde). The notably weak binding of E_8_ and K_8_ may be explained by the presence of K205 in the Class 1 pockets and E247 in the Class 2 pockets, suggesting that Class 1 pockets accommodate negatively side chains without penalty but disfavor positively charged side chains, while Class 2 pockets display the opposite binding polarity to disfavor negatively charged side chains. This explains the weak binding of homotypically charged peptides and suggests that peptides of alternative positive and negative charged side chains will bind more strongly. Consistent with this prediction, we found that (EK)_4_ and (KE)_4_ bound at least 11-67-fold tighter than E_8_ or K_8_.

The observation that (EK)_4_ binds 6-fold tighter than (KE)_4_ may reflect a preference in binding orientation. To engage favorably with all eight pockets (EK)_4_ must bind in the N to C orientation while (KE)_4_ must bind in the C to N orientation. Thus, the difference in binding affinity between (KE)_4_ and (EK)_4_ may reflect preferred N to C orientation, which is consistent with the model that Vps4p translocates from the flexible C-terminal region of ESCRT-III subunits into the folded N-terminal domains. We note, however, that this inference is far from definitive and that it is widely held that other AAA unfoldases can translocate along their substrates in either direction^19^. Regardless of this detail, our peptide-binding studies support the conclusion that the Vps4p translocation pore can bind a wide variety of amino acid sequences.

We have reported that the N-terminal region encompassing the MIT domain and linker is autoinhibitory for peptide binding, although whether this was mediated by the MIT domain or the linker (or both) was not determined^18^. We also found that Vps4p can bind circular peptides^14^, which suggested that autoinhibition might be mediated by the linker forming a hairpin conformation that plugs the translocation pore by mimicking substrate binding. To explore this possibility, we replaced the entire linker region with non-binding residues, glycine and proline, but found no effect on binding to Peptide-8 (Fig. 2a). This argues against linker-mediated autoinhibition and suggests that the effect may be a property of the MIT domain.

**Fig. 2.**
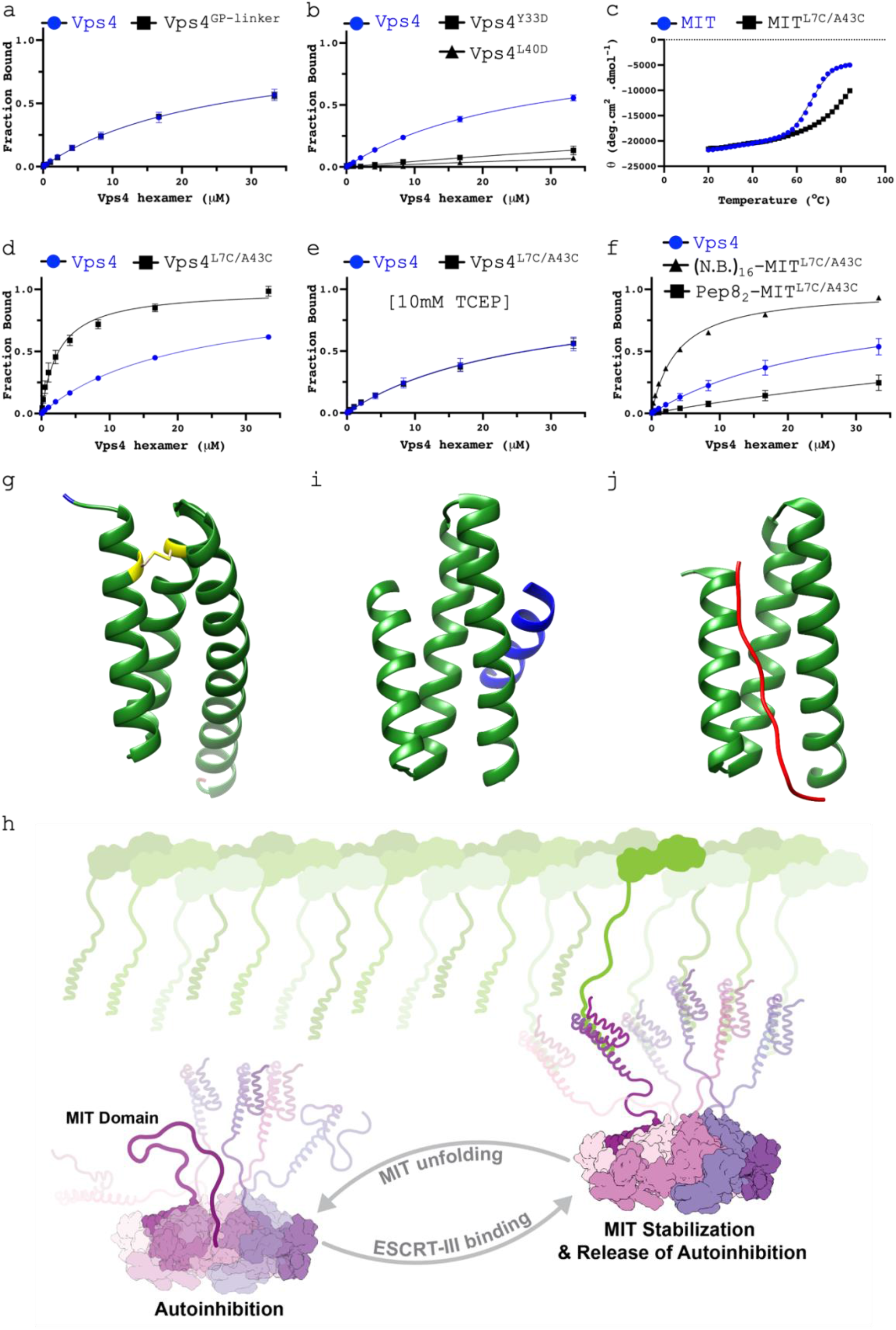
Autoinhibition involves OMIT domain unfolding. **ab**, Fluorescence polarization peptide binding assays for the indicated constructs binding to peptide-8. **c**, Melting curves (20-84°C) for MIT domains of wild type Vps4p and Vps4p^L7C/A43C^. Melting curves (20-84°C) were analyzed using the Boltzmann Sigmoidal equation to calculate Tm values, WT (66°C) and MIT^L7C/A43C^ (80°C). **def**, Binding assays as for panels ab. **g**, Vps4p MIT^L7C/A43C^ crystal structure (PDB: 9BL8). **h**, Model of autoinhibition mechanism. MIT domains fold as 3 helical bundles, but their marginal stability and proximity to the translocation pore allows unfolding to plug the pore by binding in the same manner as substrate. Binding of MIT domains to the MIM sequences displayed on ESCRT-III filaments recruits Vps4 to the ESCRT-III substrate and stabilizes MIT domains in their folded conformation away from the translocation pore. **i**, NMR structure of *H. sapiens* VPS4A MIT bound to CHMP1A (MIM1) (PDB:2JQ9)^9^. **j**, NMR structure of *H. sapiens* VPS4A MIT bound to CHMP6 (MIM2) (PDB:2K3W)^8^.

To determine if the folded structure of the MIT domain is important for autoinhibition we created two Vps4p constructs, Vps4p^Y33D^ and Vps4p^L40D^, that are each destabilized by substituting a buried MIT hydrophobic residue for aspartate. These constructs showed significant decrease in peptide-8 binding affinity, suggesting that autoinhibition involves MIT domain unfolding (Fig. 2b). The hypothesis that autoinhibition is mediated by MIT domain unfolding suggested that increasing MIT domain conformational stability would have the opposite effect. We therefore created and determined the structure of a disulfide bond-stabilized MIT^L7C/A43C^ domain that is highly resistant to unfolding (Figs. 2cg). As predicted by the hypothesis, Vps4p^L7C/A43C^ bound peptide-8 8-fold tighter than Vps4p (Fig. 2d), and this decrease in autoinhibition was reversed upon disulfide bond reduction (Fig. 2e). Therefore, the MIT domain mediates autoinhibition in an unfolded conformation, presumably by binding the substrate translocation groove in the same manner as substrate.

To further explore this mechanism, we determined the impact of adding flexible residues to the N-terminus of the MIT domain-stabilized Vps4p^L7C/A43C^ construct. One construct displayed 16 non-binding residues (GG(PGGGG)_2_PGGG; (N.B.)_16_-Vps4p^L7C/A43C^) and the other displayed two repeats of peptide-8 (Pep8_2_-Vps4p^L7C/A43C^). As expected, peptide-8 binding to Pep8_2_-Vps4p^L7C/A43C^ was almost undetectable, while binding to (N.B.)_16_-Vps4p^L7C/A43C^ was essentially identical to Vps4p^L7C/A43C^ (Fig. 2f).

In conclusion, Vps4p specificity is regulated by multiple mechanisms. Vps4 is recruited to substrate by binding of MIM sequences to the MIT domain, and the filamentous structure of ESCRT-III promotes the high local concentration needed to overcome weak Vps4p hexamerization^20^, an effect that is amplified by the cofactor protein Vta1/LIP5, which displays multiple MIT domains and binds the Vps4p hexamer^11,20^. Here, we have uncovered an additional mechanism in which the MIT domain can autoinhibit Vps4p by unfolding to mimic substrate binding in the translocation pore (Fig 2h). Since recruitment entails MIM sequences binding and hence stabilizing the folded MIT domain, this mechanism further ties recruitment to activation, and explains the curious observation that autoinhibition can be alleviated by binding of either MIM1 or MIM2 peptides ^18^, which bind to different parts of the MIT surface but in both cases will stabilize the folded conformation (Figs. 2ij)^8–10^. It will be of interest to determine the extent to which this mechanism is utilized by other MIT-containing AAA unfoldases and, more broadly, if other members of the large family of AAA unfoldases use an analogous mechanism of substrate-induced folding of an autoinhibitory sequence.

## Acknowledgements

We thank Michael Kay, Zach Cruz, James Fulcher, Judah Evangelista, and Giovanni Quichocho, for peptide synthesis and helpful discussions, and Wes Sundquist, John McCullough, Jack Dalluge, and Peter Hackett for helpful discussions. Rachel Torrez made the mechanistic model Figure 2h. This work was supported by NIH grants U54AI170856 and R01GM112080.

## Methods

### Peptide synthesis

All peptides were synthesized using solid-phase synthesis, Fmoc chemistry, and a C-terminal amide group. Unlabeled peptides were N-terminally acetylated. For peptides used in fluorescence polarization assays, 5(6)-carboxyfluorescein (Across Organics, Geel, Belgium) was coupled to the N-terminal α-amine by standard coupling conditions. All peptides had >95% purity confirmed by reverse phase (RP)-HPLC.

### Protein Expression and Purification

All constructs used (Addgene ID): Vps4p^WT^ (ID:87733), Vps4p^ΔN^ (ID:87735), Vps4p^GP-linker^ (ID:203361), Vps4p^Y33D^ (ID:203360), Vps4p^L40D^ (ID:203357), Pep8_2_-Vps4p^L7C/A43C^ (ID:203358), (N.B.)_16_-Vps4p^L7C/A43C^ (ID:203356), and the VSL domain of Vta1p (280-330) (ID:87738) were expressed in *E. coli* BL21 (DE3) RIL from vectors based on pET-151 that expressed an N-terminal 6x His-tag followed by a Tobacco Etch Virus (TEV) cleavage site. Cultures were grown in auto-induction ZY media at 37 °C until the OD600 reached (0.5-0.8) and then transferred to 21 °C for another 16 hours. Cells were harvested by centrifugation and stored at -80 °C.

All purification steps were performed at 4 °C unless otherwise stated. Frozen cell pellets were thawed at room temperature and suspended in lysis buffer (50 mM HEPES pH=7.5, 300 mM NaCl, 10 mM Imidazole). A cocktail of protease inhibitors (pepstatin A, leupeptin, aprotinin), 1 mg of DNAase, and 100 mg of lysozyme were added to the suspended cells, followed by a 30-minute incubation. The lysis solution was kept on ice during three rounds of sonication (Branson, Flat Tip), with 1 second ON and 2 seconds OFF for 2 minutes with a 5-minute rest between rounds.

Following clarification by centrifugation for 1 hour at 41,657xg, supernatant was incubated with 5 mL of Ni-NTA agarose resin for 1 hour. Resin was washed with 150 mL of lysate buffer, and His-tagged protein was eluted with elution buffer (50 mM HEPES pH=7.5, 300 mM NaCl, 500 mM Imidazole). The N-terminal His-tag was cleaved with 1 mg of TEV protease overnight during dialysis against 25 mM Tris pH=8.0, 100 mM NaCl, 1 mM EDTA, 1 mM DTT). EDTA and DTT were removed by three 2-hour rounds of buffer exchange (25 mM Tris pH=8.0, 100 mM NaCl). The solution was subsequently incubated with 5 mL of Ni-NTA agarose for 1 hour, and a gravity column used to remove His tag, uncleaved protein, and residual His-tagged TEV protease.

Cleaved protein was further purified by ion exchange chromatography on a 5 mL HiTrap Q column (Thermo Fisher Scientific) with elution over a linear gradient from 100% buffer A (25 mM Tris pH=7.5, 100 mM NaCl) to 100% buffer B (25 mM Tris pH=7.5, 1 M NaCl) over 25 column volumes. Fractions containing Vps4p constructs were concentrated and further purified by size exclusion chromatography on Superose 6 Increase 10/300 GL (Sigma-Aldrich) while MIT domains used HiPrep 16/60 Sephacryl S-100 HR (Sigma-Aldrich). Protein-containing fractions were concentrated using Vivaspin 30,000 or 3,000 MWCO spin concentrators (Sartorius), protein concentration was measured using a NanoDrop 2000c Spectrophotometer (Thermo Fisher Scientific) for subsequent experiments.

### Thermal melt CD spectroscopy

The CD spectra of the MIT^WT^ and MIT^L7C/A43C^ were obtained in triplicates using an AVIV model 410 circular dichroism spectrometer. Samples (18.5 μM protein in 10 mM phosphate, pH 7.5) and blanks (10 mM phosphate, pH 7.5) were added to a 1 mm quartz cuvette. Ellipticity change at 222 nm was measured as a function of temperature from 20 °C to 84 °C in 2 °C increments with 2 min equilibration prior to data acquisition. Reversibility was assessed by cooling from 84 °C to 20 °C by 4 °C increments with 4-minute equilibration times. All samples demonstrated appropriate refolding (>85%). The mean residue molar ellipticity was calculated following blank subtraction and plotted as a function of temperature to determine the melting temperature (T_m_). The dynode voltage at 222 nm was monitored throughout CD experiments and did not exceed 500 volts. The data were processed, and figures created using GraphPad Prism version 9.4.1 for Mac, GraphPad Software, San Diego, California USA, www.graphpad.com.

### Crystal structure determination

Crystals of MIT^L7C/A43C^ were grown by sitting drop vapor-diffusion at 21 °C with a 2:1 ratio of protein (10 mg/mL, 20 mM Tris pH=8, 100 mM NaCl) to reservoir (200 mM CaCl_2_, 100 mM Na acetate pH=4.6, 20% 2-propanol). Prior to data collection, crystals were transferred to reservoir solution diluted with 20% glycerol, suspended in a rayon loop, and plunged into liquid nitrogen.

Diffraction data were collected on beamline 12-1 of the Stanford Synchrotron Radiation Lightsource at 100 K. Data were processed using X-ray Detector Software (XDS)^21^. Phenix^21^ was used for molecular replacement using a previously determined structure of the MIT domain from *S. solfataricus* (PDB: 2V6Y) and refinement. COOT^21^ was used for model building. Only the first 100 frames of the full data set (1751 frames) were used for refinement because the disulfide bond was found to be susceptible to radiation damage when refined against data comprising the later images. Structure factors and model coordinates were deposited in the RCBS Protein Data Bank under PDB code 9BL8.

### Fluorescence Polarization Assay

Samples were incubated in binding buffer (25 mM HEPES pH 7.5, 150 mM NaCl, 10 mM MgCl_2_) at room temperature for one hour before measuring parallel and perpendicular fluorescence intensity using excitation/emission wavelengths of 485/535nm on a BioTek Synergy Neo microplate reader.

Dissociation constants (*K*_*D*_) were calculated by fitting the data with the equation FA = [Vsp4p hexamer] / (K_D_ + [Vps4p hexamer]), where FA is the normalized fluorescence anisotropy representing the “fraction bound”. The data was processed, and figures were created using GraphPad Prism version 9.4.1 for Mac, GraphPad Software, San Diego, California USA, www.graphpad.com. The error bars shown are the standard deviation from at least three independent experiments for every data point in the figure.

Low affinity fluorescence polarization assays (Figs. 1cd, 2ab, and 2e-g) used a six 1:1 dilution series from the highest concentration (200 μM Vps4p monomer and 400 μM VSL domain) with the eighth sample containing no protein. In each dilution, ADP-BeF_x_ and the fluorescently tagged peptide were held at constant concentrations of 1 mM and 60 nM, respectively. High affinity fluorescence polarization assays (Fig. 1e) used a 14 1:1 dilution series from the highest concentration of (12 μM Vps4p monomer and 24 μM VSL domain) with the 16^th^ sample containing no protein. In each dilution, ADP-BeF_x_, the fluorescently tagged peptide, and the VSL domain were held at a constant concentration, 1 mM, 0.5 nM, and 24 μM respectively.

