## Supplemental Data for "Vps4 substrate binding and coupled mechanisms of Vps4p substrate recruitment and release from autoinhibition"

### Extended Data Table 1. Data collection and refinement statistics.

| Crystal Structure | MIT (L7C/A43C) (PDB: 9BL8) |
|---|---|
| Beamline | SSRL 12-1 |
| Wavelength (Å) | 0.979460 |
| Resolution range (Å) | 25.7 - 1.3 (1.3 - 1.3) |
| Space group | P 21 21 21 |
| Unit cell (Å) | 20.7 42.9 96.5 |
| Unit cell (°) | 90 90 90 |
| Total reflections | 78080 (4676) |
| Unique reflections | 19914 (1397) |
| Multiplicity | 3.9 (3.3) |
| Completeness (%) | 90.1 (92.6) |
| Mean I/sigma(I) | 8.0 (1.7) |
| R-meas | 0.1500 (0.5555) |
| R-pim | 0.0676 (0.2675) |
| Reflections used in refinement | 19910 (1399) |
| Reflections used for R-free | 1992 (140) |
| R-work (%) | 13.60 (19.31) |
| R-free (%) | 18.03 (24.36) |
| RMS(bonds) | 0.005 |
| RMS(angles) | 0.78 |
| Ramachandran favored (%) | 100 |
| Ramachandran outliers (%) | 0 |
| Rotamer outliers (%) | 0 |
| Average B-factor | 15.48 |

Statistics for the highest-resolution shell are shown in parentheses.

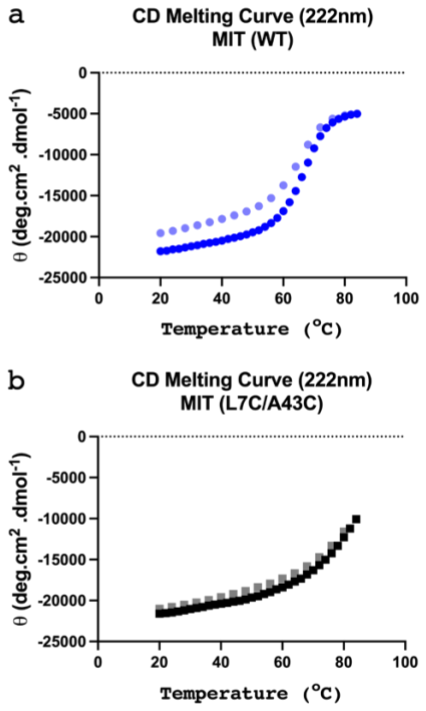


**Extended Data Fig 1. CD melting curves. a**, $MIT^{WT}$ domain melting curves from 20-84°C (blue) and from 84-20°C (light-blue) show refolding of protein at 89% of initial value. **b**, Melting curves of $MIT^{L7C/A43C}$ from 20-84°C (black) and from 84-20°C (gray) show refolding of protein at 97% of initial value.
